# A preclinical platform for focused ultrasound delivery from microplate to awake brain

**DOI:** 10.64898/2026.10.23.760846

**Authors:** Marcus S. Bell, Mark J. Williams, Chase M. Walton, Brandon W. Hughes, Terry S. Roddey, Braden B. Oldham, Claire E. Bruno, Kendell D. Peterson, B. Jacob Kendrick, Benjamin P. Caiello, Brandon A. Miller, Nathan C. Rowland, Ozgur Sahin, Ben A. Strickland

## Abstract

**Background:** Focused ultrasound (FUS) has broad therapeutic potential, but its mechanisms of action are poorly understood. Preclinical FUS research enables rigorous interrogation of understudied parameters, but fragmented, single-purpose benchtop devices prevent standardized acoustic exposure across models.

**Objective:** To describe a unified preclinical FUS platform for cell culture, ex vivo tissue, and awake stereotaxic in vivo delivery in one lightweight system, validated using glioblastoma as a disease model.

**Methods:** We performed single-well FUS delivery to 96-well microplates, single-well delivery to freshly resected human glioblastoma tissue, and stereotaxic transcranial delivery to awake, head-fixed mice ambulating freely on a floating cage. Output was independently calibrated in free field, with measured microplate insertion loss. Pulsed low-intensity focused ultrasound (pLIFU) was applied to GL261-bearing mice, resected human glioblastoma, and macrophage cell lines.

**Results:** Awake stereotaxic pLIFU produced acute tumor signal attenuation in GL261-bearing mice (mean log₂ fold-change −1.66 versus +5.20 controls, p = 0.016) with bioluminescence evidence consistent with focal targeting. Ex vivo pLIFU produced directionally consistent immunophenotypic shifts in human glioblastoma (increased iNOS+, decreased Arg1+ populations). In vitro, pLIFU reduced CD80 on immunosuppressive macrophages to 73% of baseline in RAW 264.7 cells (p = 0.0008, n = 5) and 64% in THP-1 cells (p = 0.0006, n = 2), with MHC II upregulation in RAW 264.7 (ratio 1.65, p = 0.004) and ARG1 reduction in THP-1 (ratio 0.52, p = 0.0025).

**Conclusions:** A unified preclinical FUS platform validated across in vitro, ex vivo, and awake in vivo applications enables stereotaxically guided transcranial FUS in awake, head-fixed mice ambulating in an exploratory manner during sonication, and single-well microplate treatments.

## Introduction

Focused ultrasound (FUS) is a noninvasive energy delivery technology with biological effects spanning neuromodulation to ablation.^1–3^ A FUS system comprises a signal generator, amplifier, power supply, and transducer,^4^ with target-specific adapter hardware such as transducer-laden helmets for human brain targeting, microplate scaffolds for in vitro work, and stereotaxic mounts for animal studies.^5–7^ Human transcranial systems operate at fundamental frequencies of roughly 200 kHz to 1 MHz: the Exablate Neuro system used for thalamotomy operates at 650 kHz, its low-frequency variant used for blood-brain barrier opening at 220 kHz,^55^ and human neuromodulation studies most often use 250 to 650 kHz, because lower frequencies improve skull transmission at the cost of focal resolution.^5,46^ Preclinical rodent systems operate from these frequencies up to several megahertz, where higher frequencies improve focal resolution and the thin rodent skull remains largely transmissive.^6,16^ Focusing is a property of the transducer itself, achieved by the geometric curvature of the element or by electronic beamforming in multi-element arrays; coupling to the target and dissipation of heat from the element are handled by separate subsystems (coupling media or water paths and, where needed, circulating water cooling), while the housing provides the watertight enclosure, electrical connection, and mechanical interface to target-specific hardware.^4^

FUS was developed for deep tissue ablation. Following FDA approval of MRgFUS thalamotomy for essential tremor in 2016, interest in sub-ablative low-intensity FUS (LIFU) has grown rapidly across neurology, psychiatry, and oncology. Innovation has lagged behind clinical enthusiasm because LIFU bioeffects are highly parameter-dependent and preclinical hardware is fragmented. Clinical and clinical-scale MRgFUS systems have been used productively in preclinical research, including blood-brain barrier opening and drug delivery in rat glioma models,^56,55^ nonthermal ablation in rabbits,^57^ and system evaluation in non-human primates,^58^ but such use depends on access to a clinical suite: the Insightec Exablate Neuro requires a dedicated 3T MRI scanner, over 1,300 kg of ancillary hardware, and costs ∼$2 million per device,^5^ and the focal volumes of hemispheric arrays designed for the adult human skull are better matched to rats and larger animals than to mice. Commercial preclinical systems each trade one constraint for another. MRI-guided systems^8,9^ provide treatment monitoring and thermometry but require a small-animal MRI scanner, MRI-compatible hardware, and anesthesia; stereotaxic-guided systems^10,11^ are simpler and less costly but target by atlas coordinates without image confirmation and are likewise built around anesthetized animals; and ultrasound-imaging-guided systems^12,13^ confirm transducer placement in real time but add an imaging array and the expertise to operate it. These systems cost up to $250,000 and are typically purpose-built for a single application such as mouse blood-brain barrier opening,^6^ and many rely on bulky single-element transducers with ∼50 mm apertures and motorized positioning stages^14,15^ that operate at frequencies producing focal volumes larger than the depth of the mouse brain.^16–18^ Miniaturized low-cost transducers integrated into stereotaxic frames have recently achieved sub-millimeter accuracy in mice,^6^ lowering the barrier to entry but not yet addressing delivery to cultured cells or resected tissue with the same hardware. The specific research need we address is a FUS system that works with the equipment researchers already use: a system that cannot mount on a mouse stereotaxic frame will not be adopted in neuroscience, one that cannot treat cells in a standard culture plate will not enter oncology workflows, and one that requires anesthesia cannot model human FUS, which is delivered to awake patients whose responses, including pain, can be observed. We therefore sought one acoustically characterized exposure, with a known in situ pressure, deliverable to (1) individual wells of a standard microplate with untreated controls on the same plate, (2) resected human tissue in the same format, and (3) the brain of an awake, ambulating mouse, from a single drive system at a cost accessible to a standard laboratory.^1,19^ Such a system would allow researchers to interrogate the therapeutic value, cellular mechanisms, and translatability of a hypothesized FUS parameter set with higher internal and external validity than is possible by assembling separate boutique devices for each model.

First, in vivo preclinical systems lack the precision of human FUS, which is typically an awake outpatient procedure.^20^ To our knowledge, no published platform delivers stereotaxic FUS to awake, unanesthetized mice whose heads are fixed for targeting while they ambulate omnidirectionally and at their own pace, as distinct from a linear treadmill. Additionally, current stereotaxic FUS requires anesthesia,^6^ which suppresses immune activation,^21^ alters cytokine expression,^22^ and modifies tumor growth.^23^ Implanted transducers permit awake delivery but preclude rastering and limit multi-site or multi-timepoint targeting.^24,25^ Second, translational studies of resected human tumors are limited by sonication approaches that insonate entire specimens uniformly, precluding within-specimen comparison and forcing dose-response inference across separate samples. Distributing tissue from a single resection across treated and untreated wells of one plate provides a within-patient reference that controls for inter-patient variability, time from resection, and handling, and matches the in vivo focal geometry.^26,27^ Because glioblastoma is spatially and molecularly heterogeneous,^59,60^ such a reference is imperfect: tissue sampled from different regions of the same tumor can differ in composition, a limitation we return to in the Discussion. Third, in vitro FUS experiments cannot target individual wells of a microplate while leaving adjacent wells acoustically untreated,^26–29^ so true on-plate controls and dose-response characterization without crosstalk are not yet possible. The prevailing reliance on separate culture vessels as controls introduces inter-plate variability in thermal history, media handling, and assay timing that confounds interpretation.^26^ A compact sonoporation device for 96-well plates exists but sonicates all wells simultaneously and cannot provide on-plate controls.^7^ Fourth, cost and technical complexity have limited preclinical FUS adoption. Researchers often resort to fragmented rigs from repurposed parts, which compromises internal validity and field-wide reproducibility, while off-the-shelf systems require specialized training, dedicated imaging infrastructure, and workflows unlike standard stereotaxic surgery or cell culture.^6,19^

Here we describe a benchtop FUS platform that addresses this need. It comprises a broadband RF amplifier spanning the preclinical FUS range, single-element focused transducers whose free-field output was independently calibrated, and three-dimensional acoustic field modeling that reduces reliance on empirical optimization. We use the term platform to denote the integrated drive system (signal generator, amplifier, and power supply), the control software, and the coupling adapters that deliver one calibrated exposure to a 96-well plate, a stereotaxic frame, and an awake, ambulating head-fixed mouse; the transducer itself is interchangeable, and the exposure delivered in each geometry is reported with its measured (microplate) or literature-derived (skull) transmission. For in vitro and ex vivo work, the system delivers pulsed low-intensity focused ultrasound (pLIFU) to individual wells of standard 96-well plates, permitting focal targeting of cultured cells or resected tissue with adjacent untreated controls on the same plate. For in vivo work, the platform pairs with a floating-cage head-fixation system that enables stereotaxic transcranial FUS delivery to awake, head-fixed mice ambulating freely in an exploratory manner, without anesthesia while retaining full compatibility with anesthetized stereotaxic approaches.

We selected glioblastoma (GBM) as a validation target. GBM is the most common and lethal primary brain malignancy, with median survival of approximately 15 months.^30^ Prior FUS–GBM work has focused on blood-brain barrier opening for drug delivery and liquid biopsy, with emerging interest in tumor microenvironment immunomodulation.^1,31^ GBM has not benefited from immunotherapy presumably because of its immunologically cold microenvironment and paucity of effector T cells.^32,33^ Targeting the overpopulous immunosuppressive tumor-associated macrophage (TAM) has emerged as a strategy to overcome this obstacle.^34–37^ The only prior studies showing FUS-mediated TAM reprogramming used microbubble-mediated BBB opening,^31,38,39^ where cavitation produces mechanical shear, sonoporation, and sterile inflammation distinct from pLIFU alone.^40,41^ Whether FUS acts directly on macrophages or indirectly through microenvironmental disruption remains unresolved, a question well suited to a platform delivering matched drive and pulsing parameters across awake animals, resected human tumor tissue, and macrophage monolayers. We hypothesized that identical drive and pulsing parameters delivered across in vitro, ex vivo, and in vivo modalities through one acoustically calibrated platform would produce directionally consistent biological effects in each, recognizing that in situ pressure differs with exposure geometry. We tested this in three paradigms following acoustic characterization, with matched drive and pulsing parameters throughout and geometry-specific in situ pressure estimates reported for each.

## Methods

### pLIFU platform and transducer

The FUS platform (NanoTone, Bellwether Biotech, Charleston, SC) comprises a signal generator with a maximum output of 1.0 Vpp, a 50 W broadband RF power amplifier, and a single-element, spherically focused piezoceramic element (aperture diameter 13.0 ± 0.1 mm, radius of curvature 10.0 ± 0.2 mm, f-number 0.77, resonance 1.5 MHz ± 5%, tab electrode from the concave to the convex face) of the design characterized by Hu et al.,^6^ driven at 1.5 MHz. All pressures in this manuscript are peak negative (rarefactional) pressures unless stated otherwise. The element is mounted with its back face in air in a housing 3D-printed from a thermally conductive thermoplastic polyurethane (Ice9 TPU, TCPoly, Atlanta, GA; 2 to 6 W/m·K), with the element electrodes insulated from the housing; the housing provides the air backing, the coupling face, passive heat dissipation, and the mechanical interface to the stereotaxic arm or microplate adapter, and plays no part in focusing, which is set by the curvature of the element. The housing face was coupled to the target (microplate underside or exposed skull) with ultrasound gel. For microplate work, four such transducers are held in one adapter at a 27 mm pitch, so that each addresses the center of every third well in a row. Acoustic parameters are reported following the ITRUSST consensus on standardised reporting for transcranial ultrasound stimulation,^61^ and the completed ITRUSST reporting checklist is provided as Supplementary Table 3. Two transducers of this design (T3 and T4) were mapped in-house in a degassed water tank with an automated three-axis gantry and a 0.5 mm aperture needle hydrophone (HNR-0500, Onda Corp., Sunnyvale, CA), with the transducer coupled to the tank through a window of transparent polyurethane film (Tegaderm 16004, 3M) so that the element remained air-backed as in use, following Gu et al.^49^; the film is tens of micrometers thick, a small fraction of the 1 mm wavelength at 1.5 MHz, and no correction was applied. Scans were acquired at 0.5 mm step resolution to map lateral and axial beam profiles, locate the focus, and extract peak negative pressure, peak positive pressure, pulse intensity integral, and mechanical index per ANSI/IEC standards. A k-Wave-based simulation suite^50^ generated pre-experimental field predictions that were compared with the measured beam profiles.

### Independent free-field characterization

Absolute output and focal geometry were characterized independently, on transducer T3, by Precision Acoustics Ltd (Dorchester, UK), an ISO/IEC 17025 (UKAS) accredited calibration laboratory, in degassed water with the transducer face immersed and the housing open to air (air-backed, the configuration used experimentally), driven at 1.5 MHz by the NanoTone drive system. Receivers were a 0.2 mm needle hydrophone and a 10 µm fiber-optic hydrophone, both calibrated from 1 to 30 MHz with traceability to the National Physical Laboratory (expanded uncertainty 9%, k = 2). Transverse scans at the axial pressure maximum were acquired at the drive settings giving 0.5, 1.0, and 1.5 MPa peak rarefactional pressure in water and repeated with the 96-well plate between source and receiver and the hydrophone in the target well; an axial scan provided the −6 dB focal length. Plate insertion loss was defined as IL = −20 log_10_(p_r,plate_/p_r,water_) from the spatial-peak rarefactional pressures with and without the plate. Peak compressional pressure, peak rarefactional pressure, and I_SPPA_ were measured at the spatial maximum through water and through the plate, and the maximum output of the system was measured with the fiber-optic hydrophone up to the full drive of 1.0 Vpp. Instrument details, the drive settings for each level, focal dimensions, and the maximum output are given in Supplementary Table 4.

### Derived acoustic parameters and in situ values

I_SPPA_ was computed from the measured focal waveforms by the calibration laboratory; the sinusoidal approximation I_SPPA_ = p_r_Z/(2Z) with Z = 1.5 MRayl, as used by the ITRUSST TUS calculator (https://www.itrusst.com/tus-calculator), reproduces the measured values within 6%.^61^ I_SPTA_ was obtained as I_SPPA_ × duty cycle, and the mechanical index as MI = p_r_/√f_0_ with p_r_ in MPa and f_0_ in MHz. Free-field values are reported as the primary exposure descriptor, following the ITRUSST reporting consensus.^61^ In-well values were measured directly through the plate as described above. Transcranial values in the mouse were estimated by applying the 81 ± 4% pressure transmission measured through degassed ex vivo mouse skulls at 1.5 MHz for this transducer design by Hu et al.,^6^ which agrees with the 18% attenuation reported by Choi et al.;^16^ Hu et al. also found that the mouse skull displaces the focus by less than 0.1 mm. Focal dimensions are reported at −3 dB (half of peak intensity) and −6 dB (half of peak pressure), the ITRUSST convention; the 0.44 mm region of interest used in the bioluminescence analysis lies within the −1 dB core of the measured focus (0.6 to 0.8 mm across), where pressure is within 10% of its peak. Microplate geometry was that of standard flat-bottom polystyrene 96-well plates (well bottom diameter 6.4 mm, well pitch 9.0 mm, nominal bottom thickness 1.3 mm); the −3 dB focal width (1.1 to 1.4 mm) is about one fifth and the −6 dB width (1.5 to 1.9 mm) about one quarter of the well diameter, and the −20 dB contour (2.8 mm wide) lies well inside the 5.8 mm distance from the center of the target well to the nearest wall of an adjacent well (to scale in Fig. 3d). The free-field and in-well measurements and the transcranial estimates for the three exposure levels are summarized in Table 1.

**Table 1.** Acoustic exposure parameters for the three pLIFU levels, reported per the ITRUSST consensus on standardised reporting.^61^.

| Parameter | pLIFU Low | pLIFU Mid | pLIFU High |
| --- | --- | --- | --- |
| *Free field, measured in degassed water* |  |  |  |
| Peak rarefactional pressure, $p_r$ (MPa) | 0.50 | 1.00 | 1.50 |
| Peak compressional pressure, $p_c$ (MPa) | 0.51 | 1.05 | 1.63 |
| $I_{SPPA}$ (W/cm<sup>2</sup>) | 7.85 | 33.2 | 78.0 |
| $I_{SPTA}$ (W/cm<sup>2</sup>) | 0.39 | 1.66 | 3.90 |
| Mechanical index | 0.41 | 0.82 | 1.22 |
| *In well, measured through the plate base* |  |  |  |
| Peak rarefactional pressure, $p_r$ (MPa) | 0.27 | 0.56 | 0.86 |
| Peak compressional pressure, $p_c$ (MPa) | 0.27 | 0.57 | 0.88 |
| $I_{SPPA}$ (W/cm<sup>2</sup>) | 2.23 | 9.55 | 22.7 |
| $I_{SPTA}$ (W/cm<sup>2</sup>) | 0.11 | 0.48 | 1.14 |
| Mechanical index | 0.22 | 0.46 | 0.70 |
| *Transcranial, estimated from 81% skull transmission* |  |  |  |
| Peak rarefactional pressure, $p_r$ (MPa) | 0.41 | 0.81 | 1.22 |
| $I_{SPPA}$ (W/cm<sup>2</sup>) | 5.15 | 21.8 | 51.2 |
| $I_{SPTA}$ (W/cm<sup>2</sup>) | 0.26 | 1.09 | 2.56 |
| Mechanical index | 0.33 | 0.66 | 0.99 |

### Awake, head-fixed, freely ambulating stereotaxic transcranial FUS delivery system

All in vivo pLIFU was performed in awake, head-fixed mice using a floating cage system (MouseWalk, Bellwether Biotech) that uses continuous positive airflow to support a lightweight circular cage on a near-frictionless surface, permitting omnidirectional, self-paced ambulation while the animal’s head remains rigidly fixed. This is classical head fixation combined with an air-lifted cage that the animal ambulates on and rotates beneath itself in any direction, in contrast to a linear treadmill; the head, not the animal, is stationary (Fig. 2c, d; Supplementary Videos 1 and 2). We therefore describe the preparation as awake, head-fixed, and freely ambulating in an exploratory manner, where ‘freely’ denotes unrestricted direction and pace of ambulation rather than a free head. The design extends prior air-table and floating-cage head-fixation platforms^51^ with integrated airflow and modular experimental configurations. Immediately following tumor cell injection, each mouse was fitted with a 3D-printed head ring cemented to the skull with dental acrylic (C&B Metabond) during the same surgical session. The head ring interfaces with a rigid clamp mounted above the floating cage, securing the skull relative to the stereotaxic transducer mount. Mice were habituated to head fixation in two daily 10-min sessions prior to pLIFU. For each session, the head ring was secured to the stereotaxic mount above the floating cage and the transducer was coupled to the skull via ultrasound gel. Animals remained awake throughout and ambulated in any direction within the floating cage while the head remained fixed. pLIFU was delivered transcranially through the intact skull (no craniotomy or cranial window), with the transducer positioned by stereotaxic coordinates referenced to bregma so that the geometric focus was placed at the tumor implantation site (AP −2.0, ML +2.0, DV −3.0 mm). Animals, husbandry, and tumor implantation are described under In vivo GL261 bioluminescence.

### In vivo GL261 bioluminescence

All animal procedures were approved by the Medical University of South Carolina IACUC (AUP-25-83) and conducted per NIH guidelines. Mice were housed under standard conditions on a 12 h light/dark cycle with food and water available ad libitum. Female C57BL/6J mice (8–10 weeks, The Jackson Laboratory) received intracranial implantation of 1 × 10⁵ GL-261-LUC cells (Vitro Biotech VLU3B133), a moderately immunogenic syngeneic glioma line with basal MHC I but not MHC II expression,^43,52^ at stereotaxic coordinates AP −2.0, ML +2.0, DV −3.0 mm relative to bregma under isoflurane anesthesia. Cells were maintained in DMEM with 10% FBS, 1% GlutaMAX-I, 1% HEPES (1 M), 1% penicillin-streptomycin at 37°C, 5% CO₂, with 0.5 μg/mL puromycin selection. Ten mice were allocated to two groups: pLIFU-treated (n = 5; four tumor-bearing and one sham-injected animal that received the injection procedure without tumor cells) and untreated control (n = 5; tumor-bearing). Mice were allocated by cage assignment without formal randomization, noted as a limitation. Bioluminescence from luciferase-expressing GL261-LUC cells was imaged through the intact scalp and skull with an AMI optical imaging system (Spectral Instruments): mice were anesthetized with isoflurane, injected intraperitoneally with D-luciferin (150 mg/kg), and imaged 5 min later with the standard automatic exposure of the system, without fur removal, with identical settings at baseline and 48 h after pLIFU. Post-treatment bioluminescence was measured 48 h after the single pLIFU session. Untreated controls were imaged at matched timepoints. Tumor growth was quantified as the log₂ fold-change in total emission (photons/s) from baseline to post-treatment. The sham-injected animal served as a qualitative control for pLIFU and repeated luciferin administration in the absence of tumor cells and was not included in statistical comparisons.

### Human glioblastoma ex vivo sonication

Freshly resected human glioblastoma tissue (WHO grade IV, n = 2 patients) was obtained intraoperatively and plated within one hour of resection under MUSC IRB protocol Pro00129684. Specimens were plated in a 6-well plate at 1.5 mL/well, allowed to settle 24 h at 37°C, then replated into a 96-well plate, with tissue from each specimen distributed across the treated and untreated wells so that every pLIFU condition and its control derived from the same resection, and sonicated 24 h later (approximately 48 h after resection). pLIFU was delivered in a 2 × 2 factorial design varying peak negative pressure (0.5 or 1.5 MPa) and pulse repetition frequency (5 or 25 Hz PRF; pulse duration 10 ms and 2 ms, respectively, holding the duty cycle at 5%) at 1.5 MHz for 120 s per well without microbubbles. Pressures are free-field peak negative pressures; measured in-well values are given in Table 1. Each condition was performed in duplicate wells. Cells were harvested 24 h post-sonication and analyzed by flow cytometry. TAMs were identified by sequential gating on live CD45+CD11b+ cells. Polarization was assessed by parallel quadrant gating on CD80/CD206 (surface) and iNOS/Arg1 (functional). CX3CR1 and CD163 were assessed in parallel from the CD45+CD11b+ gate. Marker expression for each pLIFU condition was normalized to the untreated control mean and expressed as fold-change. Because glioblastoma is regionally heterogeneous,^59,60^ untreated wells from the same specimen are treated as a within-patient rather than a within-region reference.

### Cell culture and immunosuppressive polarization

RAW 264.7 murine macrophages (ATCC TIB-71) were maintained in DMEM with GlutaMAX-I and 10% FBS at 37°C, 5% CO₂. THP-1 human monocytes (ATCC TIB-202) were maintained in RPMI 1640 with 10% FBS and 2 mmol/L L-glutamine under identical atmospheric conditions. THP-1 cells were differentiated to macrophages with 100 nM phorbol 12-myristate 13-acetate (PMA; Sigma P1585) for 48 h, then medium was replaced. RAW 264.7 cells were used without PMA. For immunosuppressive polarization, all wells in both cell lines were treated with 20 ng/mL recombinant human TGF-β1 (R&D Systems 7754-BH) for 24 h^53^ and left in TGF-β1 for the duration of the experiment. After 24 h, wells were spiked (2 μL into 100 μL existing medium) with either proinflammatory stimulus (LPS 100 ng/mL, Sigma L4391; plus IFN-γ 50 ng/mL, PeproTech 300-02)^53^ or an equivalent volume of vehicle. In pLIFU wells, sonication was performed concurrently with vehicle spiking in separate wells from those receiving the proinflammatory stimulus. All spiked conditions were maintained 24 h before harvest in 96-well plates with 100 μL/well.

### pLIFU treatment

Sonications used pulsed low-intensity focused ultrasound without microbubbles at 1.5 MHz center frequency, 10 ms pulse duration, 200 ms pulse repetition period (PRF 5 Hz), 5% duty cycle, and 120 s per well. Three free-field peak negative pressure amplitudes were tested, pLIFU Low (0.5 MPa), Mid (1.0 MPa), and High (1.5 MPa), as determined by in-house hydrophone calibration; on T3, the independently calibrated unit, these levels correspond to signal generator settings of 0.0645, 0.132, and 0.204 Vpp (Fig. 1b), to measured in-well peak negative pressures of 0.27, 0.56, and 0.86 MPa, and to estimated transcranial values of 0.41, 0.81, and 1.22 MPa (Table 1). For in vitro and ex vivo work, the transducer was positioned with a pointer whose tip lies at the measured focal distance: the pointer, held in the same stereotaxic arm as the transducer, was touched to the underside of the target well at its center, the transducer replaced the pointer, and the arm was advanced 1.0 mm toward the plate so that the focal peak lay at the cell monolayer through the 1.3 mm plate base; the axial extent of the focus makes this placement insensitive to small errors. The housing face was coupled to the plate underside with ultrasound gel. For in vivo work, pLIFU High (1.5 MPa free field) was delivered in a single 120 s session with the housing coupled to the exposed skull of the head-fixed mouse and the focus placed at the implantation coordinates, an exposure below the ITRUSST mechanical-index threshold for non-significant risk in the absence of microbubbles (MI ≤ 1.9).^42^

**Figure 1.**
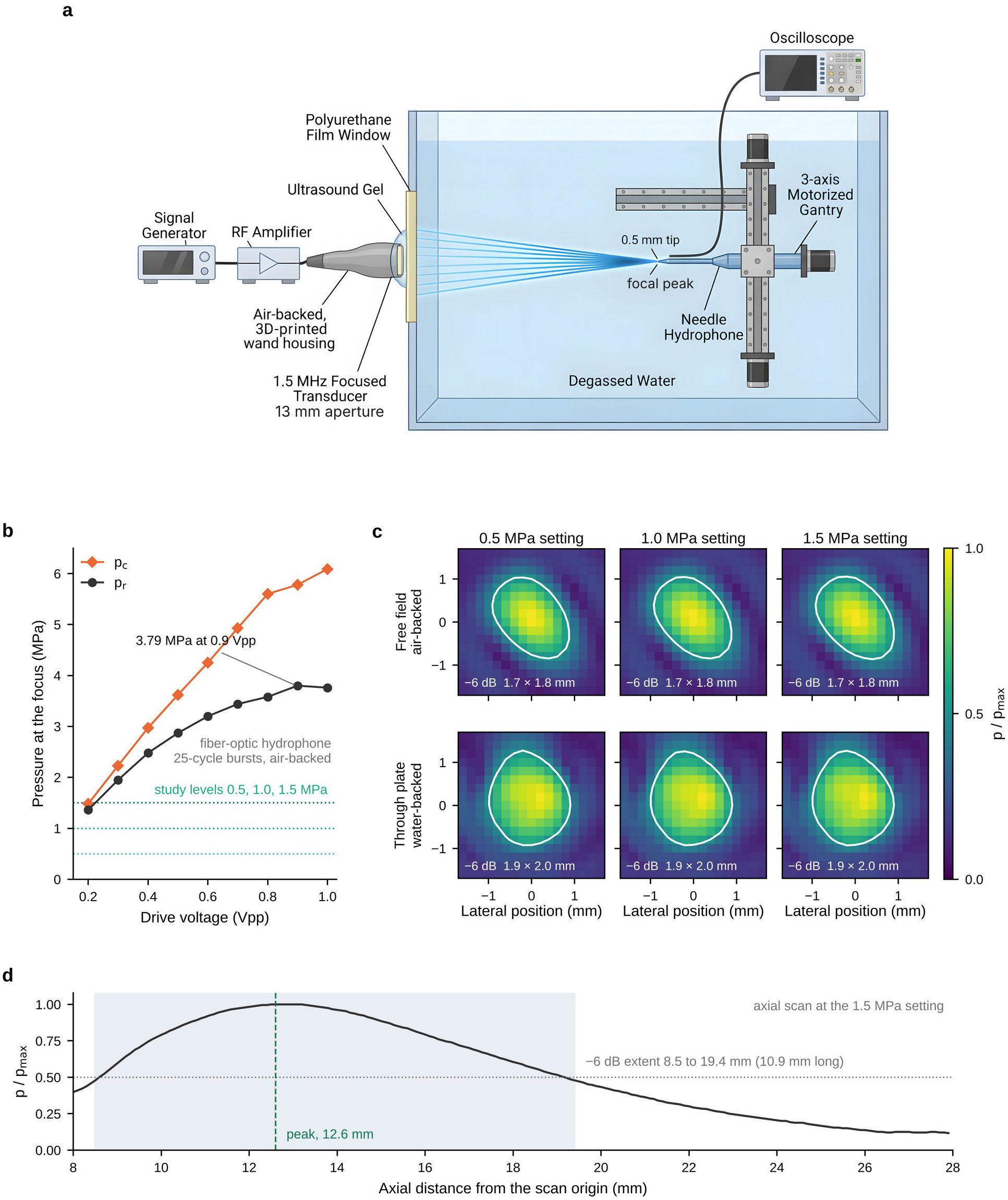
Acoustic characterization of the pLIFU platform. (a) In-house beam-mapping geometry. The air-backed transducer in its 3D-printed housing is coupled with ultrasound gel to a thin polyurethane film window in the wall of the degassed-water tank, and a 0.5 mm needle hydrophone on a three-axis motorized gantry scans the field on the far side of the window; the transducer is driven by the signal generator and RF amplifier of the drive system and the hydrophone is read on an oscilloscope. The independent characterization in b to d was performed by the calibration laboratory on transducer T3, fully submerged (Supplementary Table 4). (b) Output of T3 measured with a fiber-optic hydrophone (element air-backed, 25-cycle bursts): peak compressional (pc) and peak rarefactional (pr) pressure at the focus against drive voltage. Dotted lines mark the three exposure levels used in this study; the maximum pr was 3.79 MPa at 0.9 Vpp. At the 0.204 Vpp operating setting this survey interpolates to 1.38 MPa, against 1.50 MPa measured separately at that setting (Table 1); the 8% difference is within the expanded uncertainty of the measurement (9%, k = 2). (c) Transverse pressure maps in the focal plane measured with a 0.2 mm needle hydrophone (17 × 17 points at 0.2 mm) at the three nominal settings (0.5, 1.0 and 1.5 MPa): free field with the element air-backed (upper row) and through the base of a 96-well plate with the hydrophone inside the target well and the element water-backed (lower row). Each map is normalized to its own maximum; the white contour is the −6 dB (half-pressure) boundary and its width is given in each panel. (d) Axial pressure profile along the beam axis at the 1.5 MPa setting, normalized to the peak; the shaded band is the −6 dB extent, 8.5 to 19.4 mm from the scan origin (10.9 mm), with the peak at 12.6 mm. Absolute values for each exposure level are given in Table 1 and Supplementary Table 4.

### Parameter selection

The 1.5 MHz fundamental frequency and the transducer geometry follow Hu et al.,^6^ who showed that this element design delivers a focus of about 1.8 mm through the intact mouse skull with 81% transmission and sub-millimeter targeting accuracy, and 1.5 MHz is also the frequency used by Wintermark and colleagues for pulsed FUS in GL26 gliomas.^31,62^ Those studies used microbubbles to open the blood-brain barrier at 0.5 MPa; we omitted microbubbles, which removes the cavitation nuclei responsible for barrier opening and for the microhemorrhage seen with microbubbles,^6,40,41^ and used peak negative pressures up to 1.5 MPa, which without microbubbles remain below the ITRUSST mechanical-index threshold for non-significant risk (MI ≤ 1.9).^42^ The 10 ms pulse at 5 Hz (5% duty cycle) is a pulsed, low-duty-cycle regimen within the range reviewed for low-intensity FUS neuromodulation and immunomodulation;^44–46^ it limits the time-averaged intensity (Table 1) while delivering sustained mechanical exposure, and 25 Hz was added ex vivo to test whether pulse rate alters the response at constant duty cycle. A 120 s exposure per target (600 pulses) was chosen as a duration compatible with sonicating many wells per plate and with the length of an awake head-fixation session.

### Flow cytometry

Cells were harvested 24 h post-sonication by gentle pipetting (RAW 264.7) or Accutase incubation (THP-1), washed in FACS buffer (PBS, 2% FBS, 0.1% sodium azide), and stained for surface markers for 30 min at 4°C in the dark. For intracellular targets (ARG1, NOS2), cells were fixed and permeabilized using the eBioscience Foxp3 Staining Buffer Set (Thermo Fisher 00-5523-00) per manufacturer instructions, then stained intracellularly for 30 min at room temperature. The RAW 264.7 panel comprised MHC II, CD80, CD11b, NOS2, and ARG1. The THP-1 panel comprised HLA-DR, CD80, ARG1, CX3CR1, CD68, and CD11b. Live/dead discrimination used LIVE/DEAD Fixable Yellow (Thermo Fisher L34959). All antibodies were used at 1:100. Events were acquired on a Cytek Northern Lights spectral flow cytometer with spectral unmixing applied using single-stain controls. A minimum of 5,000 live-gate events were collected per well. Data were analyzed in FlowJo v10 (BD Biosciences). The gating strategy (debris exclusion by FSC-A vs. SSC-A, singlet selection by FSC-H vs. FSC-A, LIVE/DEAD Yellow-negative gating, then single-parameter histograms with FMO or unstained controls) was applied identically across experiments and cell lines. Percentage of live cells positive for each marker was the primary readout.

### Experimental design

RAW 264.7 experiments were performed on five separate days (n = 5 independent biological replicates); THP-1 experiments on two separate days (n = 2). Each independent experiment comprised five conditions: (1) immunosuppressive control (TGF-β1 + vehicle), (2) proinflammatory control (TGF-β1 + LPS/IFN-γ), (3) pLIFU Low, (4) pLIFU Mid, and (5) pLIFU High (each TGF-β1 + vehicle + sonication). One well per condition per experiment was used. This one-well-per-condition design treats each independent experiment as a biological replicate, the appropriate unit for between-experiment inference.^54^ Well positions were randomized per experiment, with treated and adjacent untreated wells placed in neighboring positions to confirm spatial specificity. No blinding was performed; treatment assignment was inherent to the procedure.

### Statistical analysis

All statistical analyses were performed in Python (SciPy v1.11, statsmodels v0.14, NumPy v1.24); α = 0.05 throughout. For the in vivo GL261 experiment, fold-change in total bioluminescent emission was log₂-transformed prior to analysis given right-skewed distributions. Between-group log₂ fold-change was compared by Mann-Whitney U (pre-specified, non-parametric, robust at small n) and Welch’s t-test (for completeness). Cohen’s d was calculated as the difference in group means divided by pooled standard deviation. Focal specificity was assessed by concentric ROI analysis (inner diameter 0.44 mm, within the −1 dB core of the measured focus, whose −6 dB width is 1.5 to 1.9 mm; outer diameter approximately 3-fold the inner), with total emission per ROI recorded using Aura imaging software (Spectral Instruments). The center-to-periphery ratio was defined as inner divided by peripheral (outer minus inner) emission. One mouse was excluded because complete tumor elimination yielded a mathematically undefined ratio. The change in center-to-periphery ratio was tested by one-sided paired t-test under an a priori directional hypothesis. Cohen’s d for paired data was the mean of paired differences divided by their standard deviation. For in vitro experiments, within-experiment normalization controlled for between-experiment variability in TGF-β1 polarization efficiency: each pLIFU value was divided by the same-experiment immunosuppressive control value. Normalized ratios were tested against 1.0 by one-sample t-test; raw pairwise comparisons used Welch’s t-test, and overall group differences used one-way ANOVA. Dose-response was assessed by linear regression of normalized ratio against peak negative pressure. Fifteen statistical tests were performed across the in vitro dataset; no family-wise correction was applied in the main text given the exploratory aim of identifying which markers respond to pLIFU. Benjamini-Hochberg FDR-corrected q-values are provided in Supplementary Table 2; 9 of 15 tests remained significant after correction.

## Results

### Platform characterization and acoustic calibration

In-house hydrophone mapping of two transducers of this design (T3, T4) gave linear voltage-to-pressure relationships over the drive range used experimentally (RZ > 0.99) and beam profiles that agreed with k-Wave simulations within 5% RMSE; these calibrations set the three exposure levels. Independent free-field characterization of T3 (Fig. 1b to d) established the absolute output and focal geometry of the system. Peak rarefactional pressures of 0.50, 1.00, and 1.50 MPa at the focus were obtained at signal generator settings of 0.0645, 0.132, and 0.204 Vpp, with peak compressional pressures of 0.51, 1.05, and 1.63 MPa and measured I_SPPA_ of 7.85, 33.2, and 78.0 W/cmZ (Table 1); the near-equality of compressional and rarefactional pressures shows that the experimental range lies in the linear regime. The focal width was 1.06 × 1.35 mm at −3 dB and 1.46 × 1.87 mm at −6 dB with the element water-backed (1.20 × 1.27 mm and 1.66 × 1.76 mm air-backed), did not change with drive level (within 0.05 mm across the three settings), and the −6 dB axial extent was 10.9 mm (8.5 to 19.4 mm along the beam axis, with the peak at 12.6 mm; Supplementary Table 4). With the 96-well plate in the path, the focus was preserved but broadened to 1.86 × 1.98 mm at −6 dB (Fig. 1c), and the insertion loss was 5.4 ± 0.5 dB, giving in-well peak rarefactional pressures of 0.27, 0.56, and 0.86 MPa and I_SPPA_ of 2.23, 9.55, and 22.7 W/cmZ at the three settings (Table 1). The maximum output of the system, measured with the fiber-optic hydrophone, was 3.79 MPa peak rarefactional pressure at 0.9 Vpp, more than twice the highest level used in this study (Supplementary Table 4). At the three exposure levels used in this study the free-field mechanical index was 0.41 to 1.22 and I_SPTA_ 0.39 to 3.90 W/cmZ; the in-well mechanical index was 0.22 to 0.70 and the estimated transcranial peak rarefactional pressure 0.41 to 1.22 MPa (Table 1). All exposures fall below the ITRUSST mechanical-index threshold for non-significant risk in the absence of microbubbles (MI ≤ 1.9).^42^

### Awake, freely ambulating, head-fixed stereotaxic pLIFU delivery via a floating cage system

To validate awake, head-fixed FUS delivery, we paired the platform with a floating-cage head-fixation system and delivered pLIFU transcranially to GL261-luciferase intracranial gliomas^43^ in awake, head-fixed mice ambulating freely on the floating cage. Tumor burden was assessed by bioluminescence imaging at baseline and 48 hours post-treatment (Fig. 2f). Four tumor-bearing mice received pLIFU; five additional tumor-bearing mice served as untreated controls. One sham-injected mouse (identified in Fig. 2f) also received pLIFU as a qualitative control for pLIFU and repeated luciferin administration in the absence of tumor cells; this animal was not included in statistical comparisons. All five untreated control tumors grew (median 54-fold increase, range 3- to 180-fold). FUS-treated tumors showed attenuated growth, with three of four exhibiting absolute reductions in total emission (60%, 68%, and 97% decreases in photons/s). The fourth FUS-treated mouse showed a 130% increase, substantially less than all but one control. FUS-treated tumors showed a mean log₂ fold-change of −1.66 ± 2.48 compared to +5.20 ± 2.23 for controls (Mann-Whitney U = 0, p = 0.016; Welch’s t-test, p = 0.005; Cohen’s d = −2.92; Fig. 2h). The sham-injected mouse showed only low-magnitude background signal at both timepoints (Fig. 2f).

**Figure 2.**
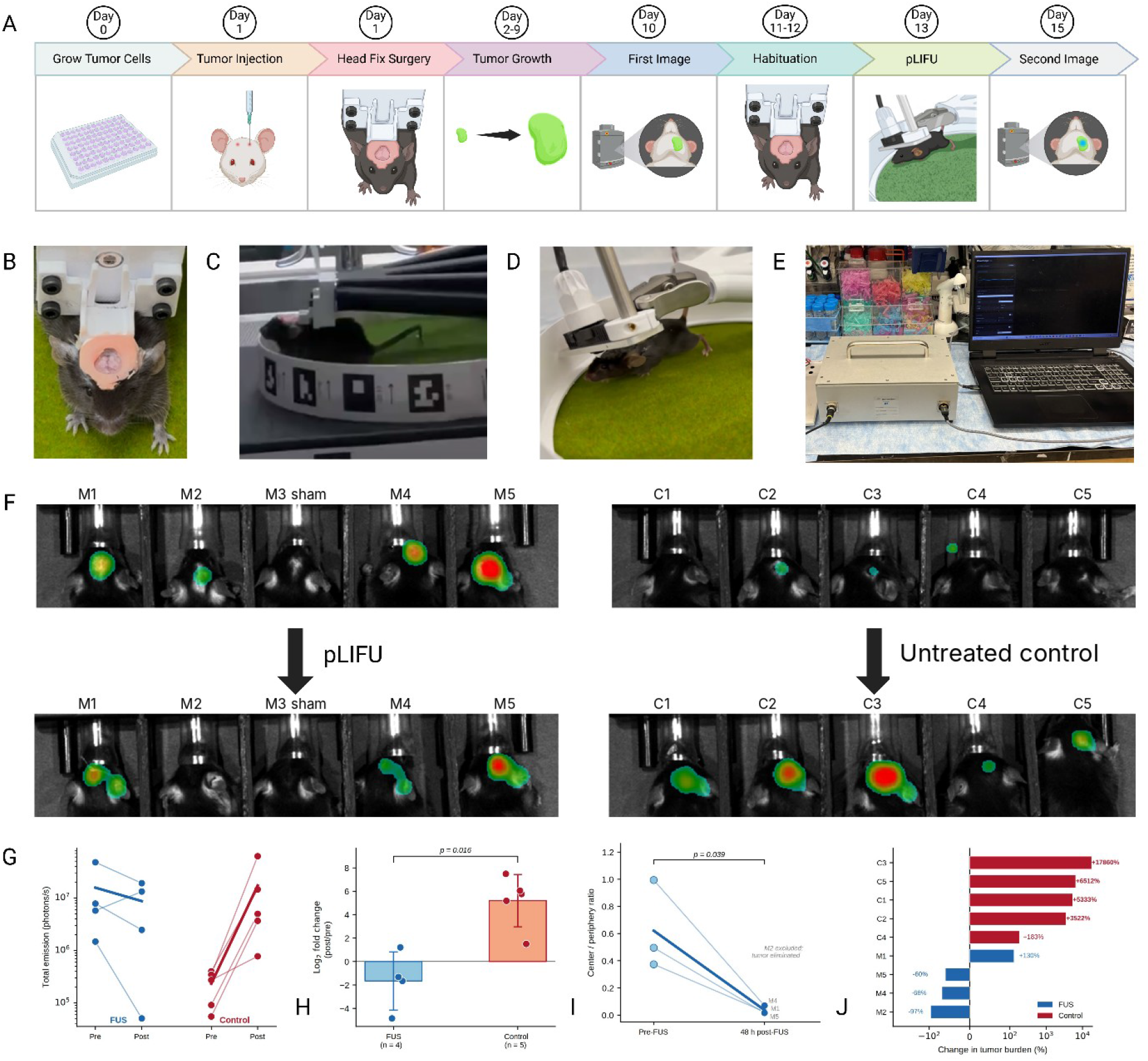
Awake stereotaxic pLIFU delivery to intracranial GL261 gliomas via a floating cage system. (A) Experimental timeline: tumor cell culture (day 0), intracranial tumor cell injection and head-ring surgery in the same session (day 1), tumor growth (days 2 to 9), baseline bioluminescence imaging (day 10), habituation to head fixation (days 11 to 12), pLIFU delivery (day 13), and post-treatment imaging (day 15, 48 h after pLIFU). (B) Mouse fitted with 3D-printed head ring cemented to the skull. (C) Mouse head-fixed above the floating cage with fiducial markers for behavioral tracking. (D) Transducer coupled to the skull surface through ultrasound gel within the head ring aperture during awake pLIFU delivery. (E) Complete FUS platform with drive system and laptop interface. (F) Bioluminescence images at baseline (upper row) and 48 h later (lower row) for the pLIFU group (left; M1 to M5, of which M3 was sham-injected without tumor cells and served as a qualitative control) and the untreated controls (right; C1 to C5, tumor-bearing). Animal numbers match panels G to J. (G) Total emission (photons/s) for individual pLIFU-treated (blue) and control (red) mice at baseline and post-treatment timepoints. (H) Log₂ fold-change in total emission for pLIFU-treated (n = 4 tumor-bearing) versus control (n = 5) groups (Mann-Whitney U = 0, p = 0.016); the sham-injected animal is not included. Box plots show median with interquartile range. (I) Center-to-periphery ratio at baseline and 48 hours post-pLIFU for three evaluable treated mice (one-sided paired t-test, p = 0.039); one mouse (M2) excluded due to near-complete signal elimination. (J) Percent change in tumor burden for individual pLIFU-treated (blue; M1, M2, M4, M5) and control (red; C1 to C5) animals; M3 (sham-injected) is not plotted. Note logarithmic scale on the x-axis.

To assess spatial targeting, we performed concentric region-of-interest (ROI) analysis. For each mouse at each timepoint, an inner circular ROI of 440 µm diameter, lying within the −1 dB core of the measured focus (0.6 to 0.8 mm across; −6 dB focal width 1.5 to 1.9 mm), was centered on the stereotaxic injection coordinates, and a larger concentric outer ROI encompassed the full tumor extent. The center-to-periphery ratio was defined as inner emission divided by peripheral annular emission. Three of four FUS-treated mice were evaluable; the fourth (97% overall reduction) was excluded because near-complete signal elimination left insufficient residual signal for spatial quantification. Among the three evaluable mice, the center-to-periphery ratio decreased from 0.621 ± 0.329 at baseline to 0.037 ± 0.029 post-FUS (one-sided paired t-test, t(2) = 3.36, p = 0.039; Cohen’s d = 1.94; Fig. 2i). Individual animal data are in Fig. 2j. One mouse whose overall tumor burden increased 130% post-FUS showed a 79% decrease in center ROI signal despite a 226% increase in annular signal, consistent with focal specificity even with net tumor progression. These results show that the floating cage system permits stereotaxically targeted transcranial pLIFU delivery in awake, head-fixed mice ambulating throughout sonication (Supplementary Video 1), with bioluminescence evidence consistent with a focal effect at the intended coordinates. The transcranial pressures at the target (Table 1) are estimated from the 81% skull transmission measured for this transducer design by Hu et al.^6^

### Single-well pLIFU delivery to cultured macrophages

To validate single-well focused ultrasound delivery in a 96-well format, we applied pLIFU to immunosuppressive macrophages (RAW 264.7 murine and THP-1 human) at three free-field pressure amplitudes (0.5, 1.0, and 1.5 MPa; measured in-well peak rarefactional pressures 0.27 to 0.86 MPa, Table 1) with untreated controls in adjacent wells of the same plate. This design tests three engineering requirements simultaneously: that the focused beam delivers acoustic energy to the target well, that adjacent wells remain acoustically untreated, and that the delivered energy produces dose-dependent biological effects.

Across all markers and conditions, CD80 reduction on pLIFU-treated immunosuppressive macrophages was the most consistent effect. In RAW 264.7 cells across five independent experiments, all three pLIFU doses significantly reduced CD80 versus immunosuppressive controls (Low p = 0.027; Mid p = 0.047; High p = 0.021, Welch’s t-test; ANOVA F = 3.64, p = 0.036). Within-experiment normalization yielded a pLIFU/immunosuppressive CD80 ratio of 0.73 (p = 0.0008, one-sample t-test vs. 1.0; Fig. 3i, k), a reduction to approximately 73% of baseline.

**Figure 3.**
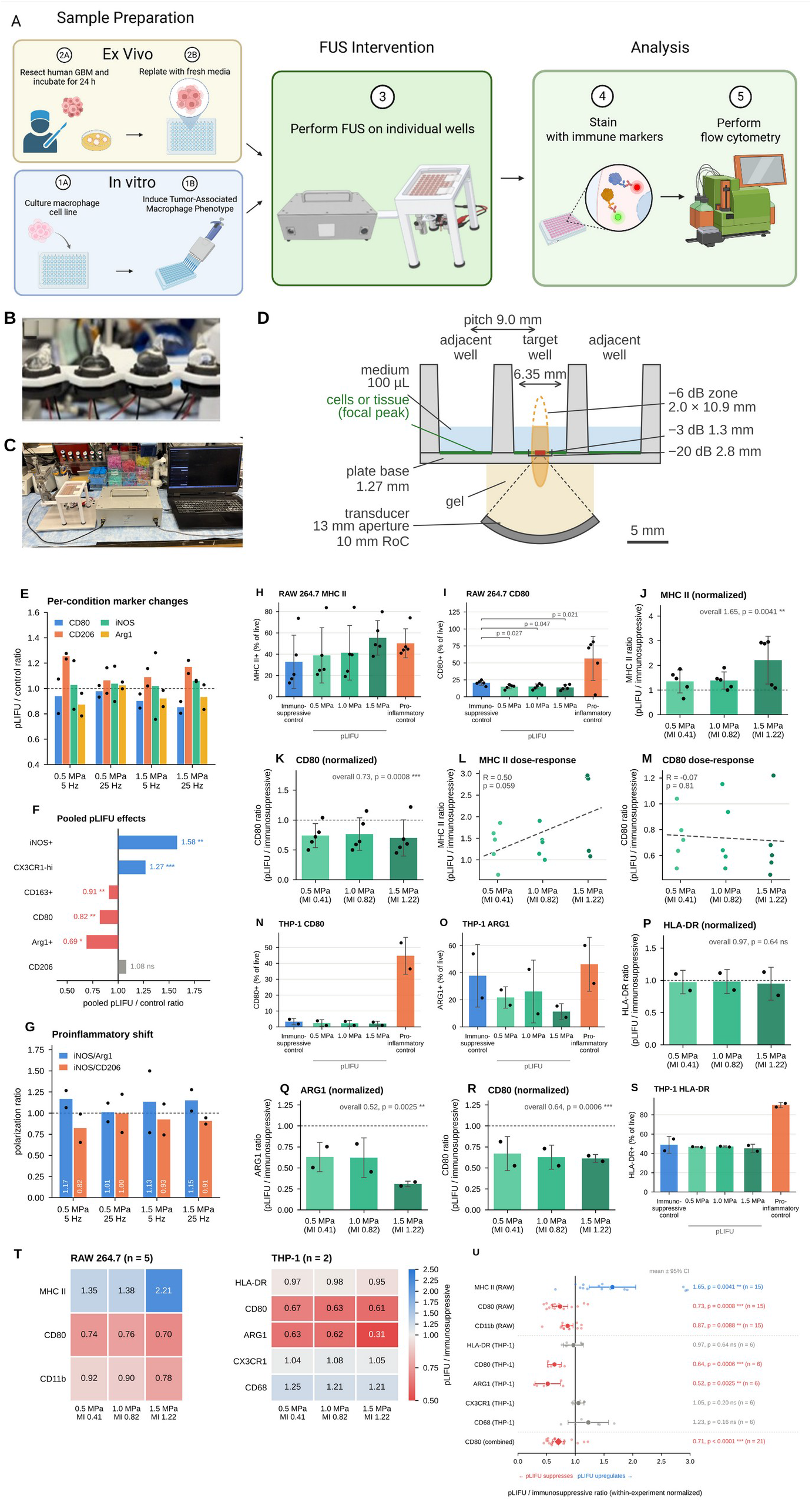
In vitro and ex vivo platform validation in immunosuppressive macrophages and human glioblastoma tissue. (A) Experimental workflows for ex vivo (upper path: resected human GBM tissue, 24-hour incubation, replating) and in vitro (lower path: macrophage culture, TGF-β1 immunosuppressive polarization) sample preparation, followed by single-well pLIFU delivery, immune marker staining, and spectral flow cytometry. (B) The four-transducer microplate adapter used for all in vitro and ex vivo delivery: air-backed transducers in sealed housings, gel-coupled to the underside of a 96-well plate at a 27 mm pitch so that each element addresses every third well; the in vivo configuration, in which a single transducer in the same housing is held in the stereotaxic arm, is shown in Fig. 2c and 2d and Supplementary Video 1. (C) FUS platform configured for in vitro delivery, with the plate seated on the adapter. (D) To-scale cross-section of the measured focus in one well of a flat-bottom 96-well plate (well bottom diameter 6.35 mm, pitch 9.0 mm, base 1.27 mm, 100 µL of medium): the orange ellipse is the −6 dB focal zone measured through the plate (2.0 mm wide, 10.9 mm long, peak placed at the cell monolayer or tissue layer; dashed above the medium surface), the red bar the −3 dB width (1.3 mm) and the bracket the −20 dB width (2.8 mm), both in free field; the single-element transducer (13 mm aperture, 10 mm radius of curvature) is gel-coupled beneath the target well. (E) Ex vivo results: pLIFU/control fold-change ratios for CD80, iNOS, CD206, and Arg1 across four pLIFU parameter combinations (0.5 or 1.5 MPa free-field peak negative pressure; 5 or 25 Hz PRF) in resected human GBM tissue (n = 2 patients; bars, mean of the two patients; points, individual patients, each the mean of duplicate wells). (F) Pooled pLIFU effects across all four parameter combinations in patient 1 (fold-change ratios relative to that patient’s untreated wells, with p-values); CX3CR1 and CD163 were measured in the same staining panel as the other markers. (G) Proinflammatory shift ratios (iNOS/Arg1 and iNOS/CD206) across ex vivo pLIFU conditions (bars, mean of the two patients; points, individual patients). (H) RAW 264.7 MHC II expression (% of live cells) across immunosuppressive control, three pLIFU doses (0.5, 1.0, and 1.5 MPa free-field peak negative pressure; MI 0.41, 0.82, and 1.22), and proinflammatory control (n = 5 independent experiments; bars, mean ± SD; points, individual experiments). (I) RAW 264.7 CD80 expression across conditions; brackets give Welch’s t-test p-values versus immunosuppressive control. (J) MHC II within-experiment pLIFU/immunosuppressive ratio by dose (overall ratio 1.65, p = 0.0041, one-sample t-test versus 1.0). (K) CD80 normalized ratio by dose (overall 0.73, p = 0.0008). (L) MHC II dose-response regression of within-experiment ratios on peak negative pressure (R = 0.50, p = 0.059). (M) CD80 dose-response regression (R = −0.07, p = 0.81). (N) THP-1 CD80 expression across conditions (n = 2 independent experiments). (O) THP-1 ARG1 expression across conditions. (P) THP-1 HLA-DR normalized ratio by dose (overall 0.97, p = 0.64). (Q) THP-1 ARG1 normalized ratio by dose (overall 0.52, p = 0.0025). (R) THP-1 CD80 normalized ratio by dose (overall 0.64, p = 0.0006). (S) THP-1 HLA-DR expression across conditions. (T) Heat maps of mean normalized ratios for all markers across doses in RAW 264.7 (left) and THP-1 (right). (U) Forest plot of pLIFU/immunosuppressive ratios (mean and 95% confidence interval, individual ratios overlaid) for all markers in both cell lines, with the combined CD80 ratio across cell lines (diamond; 0.71, p < 0.0001, n = 21). The vertical line at 1.0 indicates no effect; values below 1.0 indicate reduced expression relative to the immunosuppressive baseline. Dashed lines at 1.0 in E, G, J, K, P, Q, and R indicate no change. All pressures are free-field peak negative pressures; measured in-well values are given in Table 1.

This was replicated in THP-1 human macrophages (CD80 ratio = 0.64, p = 0.0006, n = 2; Fig. 3n, r), a 36% reduction. Pooled across species, the combined ratio was 0.71 (p < 0.0001, n = 21; Fig. 3u). RAW 264.7 cells do not constitutively express high MHC II, and immunosuppressive baseline expression varied between experiments (13–74%). Despite this, pLIFU consistently increased MHC II within each experiment (overall ratio 1.65, p = 0.004; pLIFU High 2.21, p = 0.0495; Fig. 3h, j). In early experiments with low baselines (13–21%), pLIFU High drove MHC II to 38–62%, approaching proinflammatory control levels (41–47%). In THP-1 cells, HLA-DR was unchanged (ratio 0.97, ns), possibly reflecting a ceiling effect at high PMA-induced baseline expression (immunosuppressive mean 48.9%).

In THP-1 cells, pLIFU reduced ARG1 to approximately half of immunosuppressive levels (ratio = 0.52, p = 0.0025; pLIFU High ratio = 0.31, p = 0.021; Fig. 3o, q). In RAW 264.7 cells, ARG1 expression was uniformly low (2–5%), providing insufficient dynamic range for detection. NOS2 was unchanged (ratio 1.03, ns), and macrophage identity markers (CD11b, F4/80) were largely preserved; CD11b showed a small reduction in RAW 264.7 (ratio 0.87, p = 0.009) of uncertain biological significance. No pattern consistent with classical proinflammatory polarization was observed (Fig. 3u, 3t; Supplementary Table 2); proinflammatory controls showed robust CD80 upregulation (3- to 15-fold), while pLIFU reduced CD80 below the immunosuppressive baseline.

Across seven independent experiments in two cell lines, no detectable biological effects were observed in adjacent untreated control wells, consistent with reproducible, dose-dependent biological effects through single-well focused delivery. Isolation of adjacent wells follows from the measured focal geometry (the −6 dB focal width of 1.5 to 1.9 mm is about one quarter of the 6.4 mm well diameter, and the −20 dB contour, 2.8 mm wide, lies well inside the 5.8 mm distance from the center of the target well to the nearest wall of an adjacent well) and is consistent with these biological readouts; pressure maps acquired through the plate (Fig. 1c) show that the plate attenuates and slightly broadens the focus without redirecting it. Live/dead stains were included on all cell culture flow cytometry experiments and revealed no difference between pLIFU-treated and control groups (Supplementary materials).

### Single-well pLIFU delivery to resected human glioblastoma tissue

To test delivery to resected tissue in a single-well format, we applied pLIFU to freshly resected WHO grade IV glioblastoma specimens 24 h after replating into a 96-well plate, approximately 48 h after surgical removal (n = 2 patients). Tissue from each specimen was distributed across individual wells, so that treated and untreated wells derived from the same resection, and sonicated across a 2 × 2 parameter matrix varying peak negative pressure (0.5 or 1.5 MPa) and pulse repetition frequency (5 or 25 Hz PRF) at 1.5 MHz, 5% duty cycle, without microbubbles, in duplicate wells per condition. Tumor-associated macrophages were identified by sequential gating on live CD45+CD11b+ cells and assessed by quadrant gating for CD80/CD206 (surface phenotype) and iNOS/Arg1 (functional polarization).

Pooling across all pLIFU conditions in one specimen (patient 1), pLIFU increased the iNOS+ proinflammatory population by 58% relative to untreated controls (ratio = 1.58, p = 0.002) and reduced the Arg1+ immunosuppressive population by 31% (ratio = 0.69, p = 0.027; Fig. 3f). CD80 was reduced by 18% (ratio = 0.82, p = 0.001), consistent with CD80 suppression in both RAW 264.7 and THP-1 cell lines. CD163+ cells decreased by 9% (ratio = 0.91, p = 0.002), while CX3CR1-high cells increased by 27% (ratio = 1.27, p < 0.001). These pooled statistics derive from the duplicate treated and untreated wells of that specimen; the per-condition panels show both patients (Fig. 3e, g). Per-condition analysis showed parameter-dependent variation in magnitude but not direction (Fig. 3e); CD206 immunosuppressive cells were modestly elevated across conditions. iNOS/Arg1 ratios exceeded 1.0 in all four parameter combinations (1.01 to 1.17), whereas iNOS/CD206 ratios ranged from 0.82 to 1.00 (Fig. 3g). These results provide initial evidence that the platform delivers focused ultrasound to resected human tumor tissue with sufficient energy to produce measurable immunophenotypic changes in a single-well format. Given the small number of specimens and the regional heterogeneity of glioblastoma,^59,60^ differences between treated and untreated wells may partly reflect differences in the composition of the tissue fragments allocated to each well; the mean direction of change for CD80 (decrease) and iNOS (increase) was, however, the same in all four parameter combinations (Fig. 3e), which argues against an artifact confined to a single tissue region.

## Discussion

The principal contribution of this work is a unified preclinical FUS platform which, to our knowledge, is the first to deliver acoustically calibrated FUS across cell culture, resected ex vivo tissue, and awake, head-fixed in vivo paradigms within a single, portable system. By integrating broadband amplification, calibrated transducers, 3D acoustic modeling, microplate-format single-well targeting, and a floating-cage head-fixation system, the platform addresses the research need identified above and enables standardized translational investigation that has not previously been possible.^4,19,44–46^

The validation data presented here confirm that the platform produces measurable, reproducible biological effects across all three paradigms when matched drive and pulsing parameters are applied. In awake GL261-bearing mice, pLIFU produced acute tumor signal attenuation with focal specificity at the targeted coordinates without anesthesia, demonstrating that stereotaxic transcranial FUS in awake, unanesthetized mice ambulating in an exploratory manner throughout sonication is feasible and that immunomodulatory endpoints can be studied independent of anesthesia-mediated confounds.^21–23^ In freshly resected human glioblastoma, identical pulsing parameters through the same single-well geometry produced directionally consistent immunophenotypic shifts toward an iNOS+, Arg1−, CD80-low phenotype within the live macrophage compartment. In cultured macrophages, well-specific delivery reproduced the same direction of CD80 suppression and added enhanced antigen presentation via MHC II, without adjacent-well crosstalk. The same drive and pulsing parameters produced convergent macrophage phenotypic changes across three biologically distinct preparations, although, as discussed below, the in situ pressure differed between the microplate and transcranial geometries.

The drive and pulsing parameters were identical across paradigms, and the in situ pressure differed with the coupling path: the measured insertion loss of the plate bottom (5.4 ± 0.5 dB) means that cells received 54 to 57% of the free-field peak rarefactional pressure, whereas the mouse skull transmits 81% at 1.5 MHz for this transducer design.^6^ Free-field values are therefore reported as the primary exposure descriptor, as recommended by the ITRUSST reporting consensus,^61^ together with the measured in-well and estimated transcranial values (Table 1), so that each paradigm can be compared with other work. The independent characterization also defines the range of the system: a maximum peak rarefactional pressure of 3.8 MPa, more than twice the highest level used here, and a focal zone 1.5 to 1.9 mm wide at −6 dB that fits within one well of a 96-well plate.

The glioblastoma immunomodulation model was selected as a validation target because it exemplifies a translational problem requiring rigorous multi-modal preclinical testing. Tumor-associated macrophages are the dominant immunosuppressive population in GBM, and reprogramming them is increasingly viewed as a prerequisite for immunotherapy efficacy.^37,47,48^ Whether FUS acts directly on macrophages or indirectly via microenvironmental disruption has not been resolved, in part because no prior platform could test both possibilities under matched parameters. The convergent macrophage phenotypic shifts observed here in cultured monolayers, in resected human tumor tissue, and in awake animals provide initial evidence for direct, parameter-tunable effects of pLIFU on macrophage immunobiology independent of microbubble-mediated BBB opening.^31,38,39^

Several design features merit discussion. The modular transducer interface is designed to accommodate single-element focused transducers for stereotaxic work and, in future iterations, multi-element arrays for parallel multi-well sonication; only single-element transducers were characterized here. Unlike MRI-guided systems, the platform requires no imaging suite or dedicated FUS facility and can be deployed in a standard cell culture or animal surgery suite. The floating cage system addresses the confounding effect of anesthesia on biological endpoints and permits behavioral monitoring concurrent with sonication. This opens the possibility of real-time assessment of neuromodulatory effects.

Several limitations should be noted. Transmission through the mouse skull was not measured directly in these animals; the transcranial values in Table 1 are estimates based on the 81% transmission measured for this transducer design,^6^ consistent with independent attenuation measurements at 1.5 MHz.^16^ The single sham-injected animal served as a qualitative control rather than a statistical test of pLIFU alone. The ex vivo comparisons use untreated wells from the same specimen as the reference and are subject to the regional heterogeneity of glioblastoma;^59,60^ the direction of effect was consistent across parameter combinations. The biological validation data, while consistent and directionally reproducible across paradigms, were obtained at a single early timepoint and cannot speak to durability or long-term effect. Baseline bioluminescence was higher in the treated than control in vivo group, which should be considered when interpreting fold-change comparisons. The 48-hour post-treatment window cannot distinguish durable tumor control from transient bioluminescence suppression due to edema, vascular disruption, or acute cellular stress; in vivo results should be interpreted as evidence of acute biological effect rather than sustained therapeutic efficacy. The center-to-periphery spatial analysis was performed on three evaluable animals after exclusion of the strongest overall responder and is therefore exploratory. Macrophage phenotype was characterized by flow cytometry surface and intracellular markers; transcriptomic and functional confirmation are reserved for future work. The ex vivo human cohort is small (n = 2 patients); broader replication is needed.

In summary, we have constructed and validated an integrated focused ultrasound platform that provides preclinical investigators with a single benchtop system for acoustically calibrated pLIFU delivery from cultured cells to awake, ambulating animals. The convergent biological effects observed across modalities support its utility for mechanism-driven investigation of FUS in disease models and provides a foundation for parameter optimization studies that bridge cell-level mechanisms to translational efficacy.

## Supporting information

Supplementary Video 1 - Awake pLIFU

Supplementary Video 2 - Habituation

## Acknowledgements

We thank the patients and surgical teams at the Medical University of South Carolina whose tissue donations made the ex vivo experiments possible. We thank Chris Jones and colleagues at Precision Acoustics Ltd for the independent hydrophone calibration and plate insertion-loss measurements.

## Funding

This work was supported by departmental startup funds awarded to B.A.S. by the Department of Neurosurgery, Medical University of South Carolina. The funders had no role in study design, data collection and analysis, decision to publish, or preparation of the manuscript.

## Author contributions

M.S.B. conceived the platform, designed and performed experiments, analyzed data, and led manuscript preparation. M.J.W. designed and built the drive system and transducer hardware. C.M.W. developed the platform software and contributed to manuscript revision. B.W.H. contributed to in vivo experiments. C.B. contributed to macrophage polarization methodology. K.D.P., B. J. K., and B.P.C. contributed to flow cytometry methodology. T.S.R. and B.O. contributed to manuscript revision. N.C.R, B.A.M., O.S., and B.A.S. provided mentorship, intellectual input, and critical revision of the manuscript. All authors reviewed and approved the final manuscript.

## Data availability

The data that support the findings of this study are available from the corresponding author on reasonable request. Source data for Figures 1, 2, and 3 will be provided upon publication.

## Code availability

Custom analysis scripts (Python) used for bioluminescence ROI quantification, flow cytometry post-processing, and statistical analyses are available from the corresponding author on reasonable request.

---

**Supplementary Table 1:** Flow Cytometry Antibody Panels.

RAW 264.7 (mouse) panel
| Target | Clone | Fluorochrome | Catalog No. | Supplier |
| --- | --- | --- | --- | --- |
| MHC II | NB 610-70S kit | NovaBlue 610 | M024T02B06-A | Thermo Fisher |
| CD80 | 16-10A1 | PE-Cy5 | 15-0801-82 | Thermo Fisher |
| CD11b | M1/70 | eFluor 506 | 69-0112-82 | Thermo Fisher |
| F4/80 | BM8 | SB780 | 78-4801-82 | Thermo Fisher |
| NOS2 | CXNFT | PE | 12-5920-82 | Thermo Fisher |
| ARG1 | A1EXF5 | APC | 17-3697-82 | Thermo Fisher |

THP-1 (human) panel
| Target | Clone | Fluorochrome | Catalog No. | Supplier |
| --- | --- | --- | --- | --- |
| HLA-DR | NB 610-30S kit | NovaBlue 610 | H049T03B05-A | Thermo Fisher |
| CD80 | 2D10.4 | PE-Cy5 | 15-0809-42 | Thermo Fisher |
| CD11b | ICRF44 | BV786 | 417-0118-42 | Thermo Fisher |
| CX3CR1 | 2A9-1 | AF488 | 53-6099-42 | Thermo Fisher |
| CD68 | Y1/82A | PE-Cy7 | 25-0689-42 | Thermo Fisher |
| ARG1 | A1EXF5 | APC | 17-3697-82 | Thermo Fisher |

Both panels
| Target | Clone | Fluorochrome | Catalog No. | Supplier |
| --- | --- | --- | --- | --- |
| Viability | — | Fixable Yellow | L34959 | Thermo Fisher |

**Supplementary Table 2:**
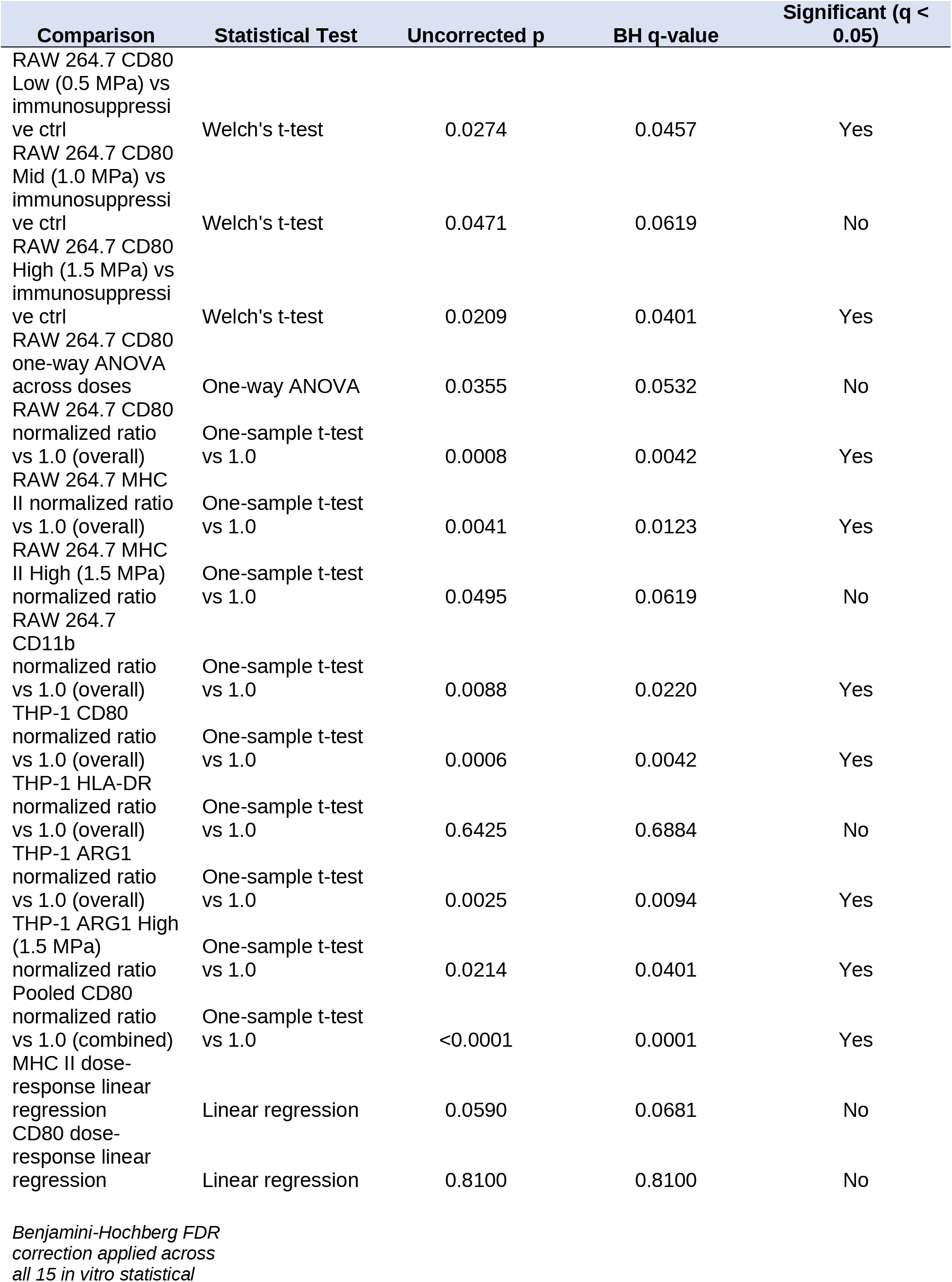
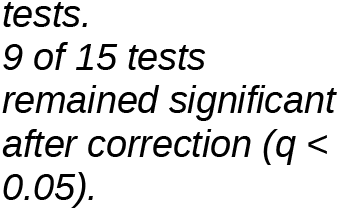
FDR.

| Comparison | Statistical Test | Uncorrected p | BH q-value | Significant (q < 0.05) |
| --- | --- | --- | --- | --- |
| RAW 264.7 CD80 Low (0.5 MPa) vs immunosuppressive ctrl | Welch's t-test | 0.0274 | 0.0457 | Yes |
| RAW 264.7 CD80 Mid (1.0 MPa) vs immunosuppressive ctrl | Welch's t-test | 0.0471 | 0.0619 | No |
| RAW 264.7 CD80 High (1.5 MPa) vs immunosuppressive ctrl | Welch's t-test | 0.0209 | 0.0401 | Yes |
| RAW 264.7 CD80 one-way ANOVA across doses | One-way ANOVA | 0.0355 | 0.0532 | No |
| RAW 264.7 CD80 normalized ratio vs 1.0 (overall) | One-sample t-test vs 1.0 | 0.0008 | 0.0042 | Yes |
| RAW 264.7 MHC II normalized ratio vs 1.0 (overall) | One-sample t-test vs 1.0 | 0.0041 | 0.0123 | Yes |
| RAW 264.7 MHC II High (1.5 MPa) normalized ratio | One-sample t-test vs 1.0 | 0.0495 | 0.0619 | No |
| RAW 264.7 CD11b normalized ratio vs 1.0 (overall) | One-sample t-test vs 1.0 | 0.0088 | 0.0220 | Yes |
| THP-1 CD80 normalized ratio vs 1.0 (overall) | One-sample t-test vs 1.0 | 0.0006 | 0.0042 | Yes |
| THP-1 HLA-DR normalized ratio vs 1.0 (overall) | One-sample t-test vs 1.0 | 0.6425 | 0.6884 | No |
| THP-1 ARG1 normalized ratio vs 1.0 (overall) | One-sample t-test vs 1.0 | 0.0025 | 0.0094 | Yes |
| THP-1 ARG1 High (1.5 MPa) normalized ratio | One-sample t-test vs 1.0 | 0.0214 | 0.0401 | Yes |
| Pooled CD80 normalized ratio vs 1.0 (combined) | One-sample t-test vs 1.0 | <0.0001 | 0.0001 | Yes |
| MHC II dose-response linear regression | Linear regression | 0.0590 | 0.0681 | No |
| CD80 dose-response linear regression | Linear regression | 0.8100 | 0.8100 | No |

*Benjamini-Hochberg FDR correction applied across all 15 in vitro statistical*
tests.
9 of 15 tests
remained significant after correction ( $q <$ 0.05).

**Supplementary Table 3.** ITRUSST reporting checklist (Martin et al., Brain Stimulation 2024, Appendix F) completed for the pLIFU exposures in this study. Free-field and microplate values are measurements by Precision Acoustics Ltd (report v1.0, 7 October 2026); transcranial values are estimates.

| ITRUSST checklist item | This study | Reported in |
| --- | --- | --- |
| **Transducer and drive system description** |  |  |
| Transducer manufacturer and model number | NanoTone focused transducer (Bellwether Biotech, Charleston, SC); single-element, spherically focused piezoceramic element assembled in-house and air-backed in a 3D-printed thermoplastic polyurethane housing | Methods, pLIFU platform and transducer |
| Transducer center frequency | 1.5 MHz (element resonance 1.5 MHz ± 5%) | Methods; Supplementary Table 4 |
| Transducer geometry | Aperture diameter 13.0 ± 0.1 mm; radius of curvature 10.0 ± 0.2 mm; f-number 0.77; single element; concave face toward the target | Methods; Supplementary Table 4 |
| Drive system components | NanoTone integrated drive system: signal generator (maximum output 1.0 Vpp), 50 W broadband RF power amplifier, power supply, and control software (Bellwether Biotech) | Methods |
| **Drive system settings** |  |  |
| Operating frequency | 1.5 MHz | Methods; Table 1 |
| Output level settings | Signal generator 0.0645, 0.132, and 0.204 Vpp, giving free-field peak negative pressures of 0.50, 1.00, and 1.50 MPa (pLIFU Low, Mid, High) on transducer T3, the independently calibrated unit | Methods; Supplementary Table 4 |
| Focal position settings | In vitro and ex vivo: a pointer whose tip sits at the measured focal distance is touched to the underside of the target well at its center, the transducer replaces the pointer in the same stereotaxic mount, and the mount is advanced 1.0 mm toward the plate so that the focal peak lies at the cell monolayer through the 1.3 mm plate base. In vivo: transducer positioned by stereotaxic coordinates relative to bregma with the focus at the implantation coordinates (AP -2.0, ML +2.0, DV -3.0 mm); housing face coupled to the exposed skull with ultrasound gel | Methods, pLIFU treatment |
| Description of transducer coupling method | Ultrasound gel between the housing face and the underside of the 96-well plate (in vitro, ex vivo) or the exposed skull within the head ring (in vivo); no cranial window | Methods |
| **Free-field acoustic parameters** |  |  |
| Reference position for measurements | Axial positions are given along the beam axis from the scan origin of the calibration laboratory (UMS4 scanning tank, Precision Acoustics); transverse scans were taken in the plane of the axial pressure maximum | Methods, Independent free-field characterization; Supplementary Table 4 |
| Spatial-peak pressure amplitude | Peak negative pressure 0.50, 1.00, 1.50 MPa; peak compressional pressure 0.51, 1.05, 1.63 MPa (0.2 mm needle hydrophone, 25-cycle bursts, air-backed configuration) | Results; Table 1 |
| Position of spatial-peak pressure amplitude | 12.6 mm from the reference position (axial scan at 1.5 MPa) | Results; Table 1 |
| Size of focal volume (-3 dB and -6 dB axial lengths and lateral widths) | Lateral -3 dB: 1.06 × 1.35 mm (water-backed), 1.20 × 1.27 mm (air-backed). Lateral -6 dB: 1.46 × 1.87 mm (water-backed), 1.66 × 1.76 mm (air-backed); unchanged across 0.5, | Results; Supplementary Table 4 |
| | 1.0 and 1.5 MPa. Axial -6 dB: 10.9 mm (8.5 to 19.4 mm from the scan origin, peak at 12.6 mm); axial -3 dB: approximately 7.4 mm, read from the same axial profile (Fig. 1d). Through the plate, lateral -6 dB $1.86 \times 1.98$ mm (-6 dB area $3.1 \text{ mm}^2$ versus $2.3 \text{ mm}^2$ in water) | |
| Position of center of focal volume | 12.6 mm from the reference position (-6 dB center) | Results; Table 1 |
| Description of how free-field parameters were obtained | Precision Acoustics Ltd (Dorchester, UK), ISO/IEC 17025 (UKAS) accredited; 0.2 mm needle hydrophone NH0200 (SN 4351) with HP33673 preamplifier (31 and 43 mV/MPa at 1 and 2 MHz, $\pm 9\%$, $k = 2$, calibrated 1 to 30 MHz, NPL-traceable) and 10 $\mu\text{m}$ fiber-optic hydrophone FOHS237/FSV2-6801-13 (137 and 157 mV/MPa, $\pm 9\%$); filtered, deionized, degassed water at 20 °C; $17 \times 17$ point transverse scans at 0.2 mm; transducer driven by the NanoTone drive system. In-house beam mapping with a 0.5 mm needle hydrophone (Onda HNR-0500) at 0.5 mm step, transducer coupled through a polyurethane film window to keep the element air-backed | Methods; Supplementary Table 4 |
| **Pulse timing parameters** |  |  |
| Pulse timing table | Pulse duration 10 ms (2 ms for the 25 Hz ex vivo condition); pulse repetition interval 200 ms (40 ms); pulse repetition frequency 5 Hz (25 Hz); duty cycle 5%; rectangular pulses, no ramping; pulse train duration 120 s (600 pulses at 5 Hz); one pulse train per well or target; a single 120 s session per animal in vivo | Methods; Table 1 |
| **In situ estimates of exposure parameters** |  |  |
| Estimated in situ spatial-peak pressure amplitude | Microplate, measured through the plate with the hydrophone realigned to the spatial peak: peak negative 0.27, 0.56, 0.86 MPa; peak compressional 0.27, 0.57, 0.88 MPa (insertion loss $5.4 \pm 0.5$ dB). Transcranial, estimated: 0.41, 0.81, 1.22 MPa using the $81 \pm 4\%$ pressure transmission measured through degassed ex vivo mouse skulls for the same transducer design at 1.5 MHz (Hu et al. 2022), consistent with the 18% attenuation reported by Choi et al. 2007 | Results; Table 1 |
| Estimated in situ pressure amplitude at the target | As above; the focal peak is placed at the monolayer (in vitro) or the implantation coordinates (in vivo) | Table 1 |
| Estimated in situ mechanical index | Microplate 0.22, 0.46, 0.70; transcranial 0.33, 0.66, 0.99 (free field 0.41, 0.82, 1.22); all below the ITRUSST threshold of 1.9 | Table 1 |
| Thermal metric (temperature rise, thermal index, or thermal dose) | Not measured. First-order estimate for brain at pLIFU High: estimated in situ ISPTA $2.56 \text{ W/cm}^2$ and an absorption coefficient of 0.9 dB/cm at 1.5 MHz give a volumetric heating rate of $0.5 \text{ W/cm}^3$ within the -6 dB focal cylinder (radius 0.85 mm); with tissue conductivity $0.5 \text{ W/(m}\cdot\text{K)}$, no perfusion, and conduction only, the steady-state temperature rise on the beam axis is approximately 0.2 °C, below the ITRUSST 2 °C limit. Heating of the culture medium is negligible (water absorption $0.005 \text{ dB/cm}$ at 1.5 MHz); absorption in the plate base was not modelled | Supplementary Table 3 |
| Description of how in situ estimates were obtained | Microplate: direct hydrophone measurement through the plate (Precision Acoustics). Skull: free-field values derated by the 81% pressure transmission measured through ex vivo mouse skull for the identical transducer design at 1.5 MHz (Hu et al. 2022) | Methods |
| **Intensity parameters (optional)** |  |  |
| Spatial-peak pulse-average intensity | Free field 7.85, 33.2, $78.0 \text{ W/cm}^2$ (computed from measured waveforms by the calibration laboratory); microplate, | Results; Table 1 |
|  | measured, 2.23, 9.55, 22.7 W/cm<sup>2</sup>; transcranial, estimated, 5.15, 21.8, 51.2 W/cm<sup>2</sup> |  |
| Spatial-peak time-average intensities | Free field 0.39, 1.66, 3.90 W/cm<sup>2</sup>; microplate 0.11, 0.48, 1.14 W/cm<sup>2</sup>; transcranial 0.26, 1.09, 2.56 W/cm<sup>2</sup> (ISPPA × 5% duty cycle, per pulse train) | Table 1 |
| Acoustic impedance used for the conversion | Measured waveforms processed by the calibration laboratory; the sinusoidal approximation with $Z = 1.5 \text{ MRayl}$ reproduces the measured ISPPA within 6% | Methods |

**Supplementary Table 4.** Transducer, drive settings, instrumentation, focal geometry, and maximum output of the pLIFU system. Free-field and through-plate values are from the independent characterization by Precision Acoustics Ltd (measurement report v1.0, 7 October 2026, and hydrophone calibration certificates); the per-level exposure values derived from these measurements are in Table 1 of the main text. pr, peak rarefactional pressure; IL, insertion loss.

| Item | Value |
| --- | --- |
| **Transducer and drive** |  |
| Transducer | Single-element, spherically focused piezoceramic element; aperture diameter $13.0 \pm 0.1$ mm; radius of curvature $10.0 \pm 0.2$ mm; f-number 0.77; resonance $1.5$ MHz $\pm 5\%$; mounted air-backed in a sealed 3D-printed housing of thermally conductive thermoplastic polyurethane that provides the coupling face, passive heat dissipation, and the mechanical interface |
| Drive system | NanoTone drive system (Bellwether Biotech, Charleston, SC): signal generator with a maximum output of 1.0 Vpp, 50 W broadband RF power amplifier, and control software; operated at 1.5 MHz |
| Signal generator setting for each level (transducer T3, independently calibrated) | pLIFU Low 0.0645 Vpp; pLIFU Mid 0.132 Vpp; pLIFU High 0.204 Vpp, giving free-field peak rarefactional pressures of 0.50, 1.00, and 1.50 MPa |
| Microplate adapter | Four transducers at a 27 mm pitch, each addressing the center of every third well of a 96-well plate (well pitch 9.0 mm); housing face coupled to the plate underside with ultrasound gel |
| Exposure used in vivo | pLIFU High (1.50 MPa free field; 1.22 MPa estimated transcranial), one 120 s session with the transducer at the implantation coordinates |
| **Independent characterization (Precision Acoustics Ltd, Dorchester, UK; ISO/IEC 17025, UKAS)** |  |
| Measurement conditions | UMS4 scanning tank; filtered, deionized, degassed water at 20 °C; transducer fully submerged and air-backed; tone bursts at 1.5 MHz from the NanoTone drive system (25 cycles for the output survey and for the measurements at the three operating settings); transducer T3; transverse scans of $17 \times 17$ points at 0.2 mm steps in the plane of the axial pressure maximum, through water and with a 96-well plate between source and receiver (hydrophone in the target well) |
| Hydrophones | 0.2 mm needle hydrophone NH0200 (serial 4351) with HP33673 preamplifier: 31 and 43 mV/MPa at 1 and 2 MHz. 10 μm fiber-optic hydrophone FOHS237 with sensor FSV2-6801-13: 137 and 157 mV/MPa at 1 and 2 MHz. Both calibrated from 1 to 30 MHz with traceability to the National Physical Laboratory; expanded uncertainty 9% ($k = 2$) |
| Free-field focal dimensions | −3 dB lateral width $1.06 \times 1.35$ mm (water-backed), $1.20 \times 1.27$ mm (air-backed); −6 dB lateral width $1.46 \times 1.87$ mm (water-backed), $1.66 \times 1.76$ mm (air-backed); unchanged across the 0.5, 1.0, and 1.5 MPa settings (within 0.05 mm). −6 dB axial length 10.9 mm (8.5 to 19.4 mm from the scan origin; pressure maximum at 12.6 mm) |
| Focal dimensions through the plate base | −6 dB lateral width $1.86 \times 1.98$ mm (water-backed); −6 dB area 3.1 mm<sup>2</sup> versus 2.3 mm<sup>2</sup> in water |
| Plate insertion loss | $5.4 \pm 0.5$ dB (mean ± SD of six measurements): 4.95 to 5.03 dB water-backed, 5.63 to 5.86 dB air-backed; $IL = -20 \log_{10}(pr,plate / pr,water)$ from the spatial-peak rarefactional pressures with and without the plate, with no axial repositioning |
| Maximum output | Peak rarefactional pressure 3.79 MPa at 0.9 Vpp, measured with the fiber-optic hydrophone (25-cycle bursts; the value at 1.0 Vpp was not higher). Needle-hydrophone data extrapolated to 1.0 Vpp give 4.15 MPa (water-backed) and 4.36 MPa (air-backed) |
| **In-house beam mapping** |  |
| Scanning system | Degassed water tank with an automated three-axis gantry and a 0.5 mm needle hydrophone (HNR-0500, Onda Corp., Sunnyvale, CA); 0.5 mm step; transducer coupled to the tank through a transparent polyurethane film window so that the element remained air-backed as in use |
| Use of the in-house data | Lateral and axial beam profiles, focus location, and comparison with k-Wave simulations (Methods); absolute pressures, intensities, and focal dimensions are taken from the independent characterization above |

**Supplementary Video 1.**
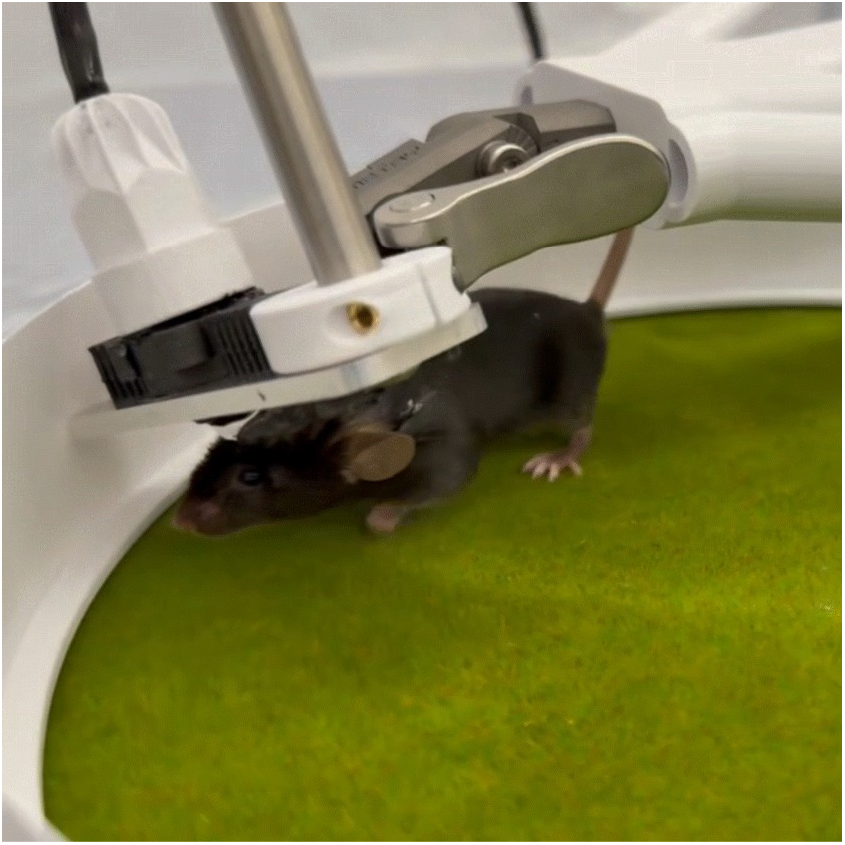
Awake, ambulating, stereotaxically guided pLIFU (MP4, 8 s, no audio). Close-up of a head-fixed mouse after habituation, with the 3D-printed head ring clamped in the head-fixation mount and the transducer in its housing coupled to the skull within the ring during pLIFU delivery, stepping on the air-lifted floating cage. The clip is 2.8 s at real time, shown three times in succession. The head is stationary and the animal ambulates; delivery is transcranial through the intact skull. The still above is a frame from the clip.

**Supplementary Video 2.**
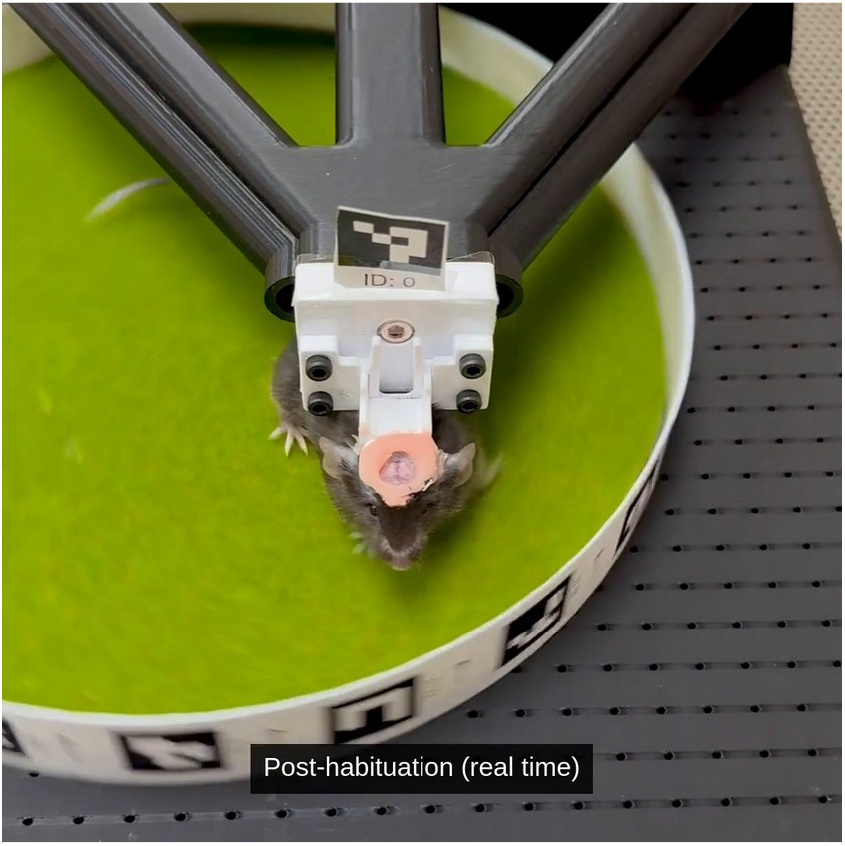
Habituation to head fixation on the floating cage (MP4, 33 s, no audio, real time throughout). Habituation day 1 (side view) and day 2 (overhead view): a head-fixed mouse in the clamp above the air-lifted cage, which it rotates beneath itself as it ambulates. Post-habituation (front view): the animal ambulating in the head-fixation clamp. The fiducial markers on the cage rim and on the mount are used for behavioral tracking. The still above is a frame from the post-habituation segment.

## Notes

### Competing Interest Statement

M.S.B., M.J.W., and C.M.W. are co-founders and equity holders of Bellwether Biotech LLC, the company that manufactures the focused ultrasound platform and floating-cage head-fixation system described in this manuscript. M.S.B. serves as Chief Executive Officer of Bellwether Biotech LLC. M.S.B., M.J.W., and C.M.W. are also named inventors on provisional patent applications related to the technology described herein, with M.S.B. as primary inventor. These applications are assigned to Bellwether Biotech LLC. Bellwether Biotech LLC has received seed-stage investment that supported development of the platform. The remaining authors declare no competing interests.

